# Low-Cost Solution for Rodent Home-Cage Behaviour Monitoring

**DOI:** 10.1101/342501

**Authors:** Surjeet Singh, Edgar Bermudez Contreras, Mojtaba Nazari, Robert J. Sutherland, Majid H. Mohajerani

## Abstract

In the current research on measuring complex behaviours/phenotyping in rodents, most of the experimental design requires the experimenter to remove the animal from its home-cage environment and place it in an unfamiliar apparatus (novel environment). This interaction may influence behaviour, general well-being, and the metabolism of the animal, affecting the phenotypic outcome even if the data collection method is automated. Most of the commercially available solutions for home-cage monitoring are expensive and usually lack the flexibility to be incorporated with existing home-cages. Here we present a low-cost solution for monitoring home-cage behaviour of rodents that can be easily incorporated to practically any available rodent home-cage. To demonstrate the use of our system, we reliably predict the sleep/wake state of mice in their home-cage using only video. We validate these results using hippocampal local field potential (LFP) and electromyography (EMG) data. Our approach provides a low-cost flexible methodology for high-throughput studies of sleep, circadian rhythm and rodent behaviour with minimal experimenter interference.

## Introduction

Rodents have been widely used for understanding the aetiology, pathophysiology and pharmacology of neurological disorders; due in part to its similar brain architecture to primates and complex behavioural repertoire. Recent improvement of sophisticated tools used for measuring and manipulating brain activity, including optogenetics, two-photon imaging, wide-field mesoscale imaging, fibre photometry, mini endoscopes [1–7], as well as the availability of diverse transgenic lines and disease models, further incentivize their use in mechanistic studies [8–11]. Though non-human primates have been traditionally used to understand the neural mechanisms underlying executive function and higher-order cognition (such as decision making, motor skill execution, and perceptual discrimination), high-throughput experiments on non-human primates are not feasible because of cost and regulation constrains [12].

The organization of spontaneous home-cage behaviour of rodents reflects the interplay between multiple neurobiological processes and can provide a high-throughput phenotyping strategy without experimenter’s intervention. In addition to the lack of observer bias in monitoring animal behaviour [13], one major advantage of automated home-cage monitoring is continuous and accurate monitoring particularly during dark periods when mice are most active [14, 15]. Furthermore, remote monitoring of animals also greatly reduces the requirement for animal handling which may be stressful and/or confound studies [15–18]. In certain disease models assessing the signs of ill health, pain and distress is difficult [19] and since mice are crepuscular and nocturnal animals making observations at the wrong time of day can result in missing them altogether. Remote home-cage monitoring systems will allow scientists, veterinarians and animal technicians to monitor animal well-being even during the dark phases [20].

There are multiple pre-existing methods currently used to evaluate animal activity in the home-cage with different degrees of invasiveness and precision. Each of these approaches have their own advantages and pitfalls. In mice, subcutaneously implanted small magnets have been used to determine animal activity with respect to an array of magnetic field sensors under the home-cage [21]. Flores *et al*. developed a non-invasive approach for monitoring animal activity by using piezoelectric sensors positioned on the cage floor [22]. Another non-invasive system to quantify sleep-awake cycles in mice has been developed using passive infrared (PIR) motion sensors [23]. Similarly, Radiofrequency identification (RFID) tags have been used to track multiple animals to study their activity in the home-cage. However, this method has the disadvantage of being expensive, having poor spatial and temporal resolution as well as the invasiveness of the tag implantation which may provide discomfort to the animal [24]. Approaches with minimal invasiveness include running wheels and video-based methodologies. Though running wheels are widely used because of their low cost and ease of implementation for measuring the rodent activity and/or changes in circadian rhythm, they have also been shown to greatly affect animal behaviour.

Electroencephalogram (EEG) and electromyogram (EMG) signals have been used as the standard approach to classify sleep-wake states, but this involves the implantation of electrodes for recordings [25]. Not only is this approach time-consuming and invasive but also expensive to implement, particularly in high throughput experiments for assessing sleep-wake behaviour under varying pharmacological and environmental manipulations [26]. In the past, several attempts have been made to develop alternative less invasive methods (other than EEG/EMG) to classify sleep-awake behaviour in rodents. For instance, video monitoring has been used to evaluate periods of sustained immobility as a surrogate of EEG/EMG-defined sleep [27, 28]. All these methods have been able to provide 90-95% agreement with tethered EEG/EMG recordings, however these methods are sometimes difficult to implement because of their technical complexity or the expense of the required software/hardware which makes them unsuitable for high throughput studies of animal behaviour. Here, we present a low-cost rodent home-cage behaviour monitoring system built around the Raspberry Pi (RPi) using open-source software technologies that is suitable for high-throughput data acquisition with virtually no experimenter interventions. We show that our system is capable to reliably video-monitor multiple home-cages simultaneously and remotely at variable frame rates (from 10-120 Hz). To demonstrate the capabilities of our system, we present a reliable sleep-wake classification based only on video data collected in the home-cage and validate it against a standard EEG/EMG based scoring.

In summary, low-cost automated home-cage monitoring and analysis systems, like the one proposed here, represent a platform upon which custom systems can be built, allowing researchers to study rodent behaviour with minimal experimenter interference [29–32].

## Material and Methods

### Animals

All experiments were carried out on adult (20-30 g, age 4 month) wild type C57BL/6 mice (n = 10). Mice were housed under standard conditions, in clear plastic cages under 12 h light, 12 h dark cycles. All the animals were either grouped or singly housed in Optimice® (Animal Care Systems) home-cages depending on the experimental protocol they were going through. Mice were given ad libitum access to water and standard laboratory mouse diet. All protocols and procedures were approved by the Animal Welfare Committee of the University of Lethbridge and were in accordance with guidelines set forth by the Canadian Council for Animal Care.

### Surgery

Local field potential (LFP) and electromyography (EMG) electrodes were implanted in C57BL6J mice anesthetized with isoflurane (2.5% induction, 1-1.5 maintenance). For hippocampal LFP recordings, a monopolar electrode made of Teflon coated stainless-steel wire (bare diameter 50.8 µm) was inserted in the brain (at coordinates -2.5mm AP, 2mm ML and 1.1mm DV from bregma), targeting the CA1 region. For EMG recordings, a multi-stranded Teflon-coated stainless-steel wire (bare diameter 127 µm) was inserted into the neck musculature using a 23-gauge needle. The reference and ground screws were placed on the skull over the cerebellum. The other end of electrode wires were clamped between two receptacle connectors (Mill-Max Mfg. Corp.) glued to the headpiece which in turn was secured to the skull using metabond and dental cement.

### Electrophysiology

After electrode implantation, the animals were allowed to recover for 7 days before being habituated to the recording setup consisting of a home-cage without the lid placed in the middle of a large Plexiglas box used as secondary containment. During 24 hour LFP and EMG recordings, the electrophysiological tethers were connected to a motorized commutator (NeuroTek Inc.). The hippocampal LFP and neck EMG signals were amplified, filtered (0.1-2000 Hz) and digitized at 4 kHz using a Digital Lynx SX System (Neuralynx, Inc.) and Cheetah software (Neuralynx, Inc.).

### System architecture/Hardware

Our system consists of a RaspberryPi (model 3B+), an infrared Picamera module (v2 NoIR), a wide-angle lens (AUKEY optics) and IR LEDs (DC12V SMD3528-300-IR (850nm)) placed on the home-cage lid to monitor the mouse behaviour (Fig. 1). To make sure that the field of view covered the entire home-cage, we placed the camera lens on the home-cage lid through a custom made hole at its centroid and sealed it with hot glue. The Picamera module is connected to the Mobile Industry Processor Interface (MIPI) Camera Serial Interface (CSI) port on the RaspberryPi via a 15cm ribbon cable. The RaspberryPi further connects to the computer network either via a LAN cable or a Wi-Fi connection (Fig. 1 A-D).

**Figure 1.**
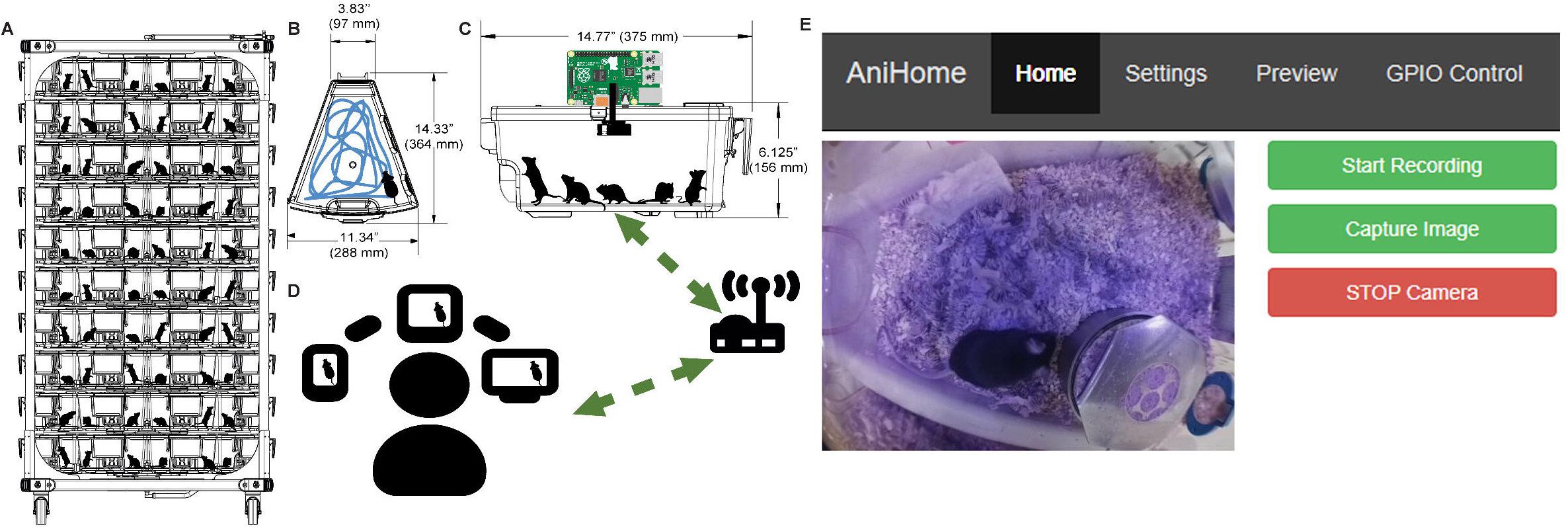
Animal home-cage monitoring system architecture and web interface. (A) Basic rack with multiple home-cages. (B) Top view and, (C) side view of a mouse home-cage with a wide-angle lens and camera mounted on top and connected to a Raspberry Pi (RPi). This RPi is responsible for data acquisition and for controlling external peripherals. The RPi further connects to a computer network either via Ethernet or Wi-Fi connection, (D) allowing the remote user to view the animal activity in the home-cage, via web browser. (E) The home page of the web interface allows the user to view the animal activity in its home-cage in real-time and to start/stop recording videos and to capture images.

### System interface

The remote user can view the live feed of the animal activity in the home-cage via a customized web interface (Fig. 1 E). In addition to live home-cage monitoring, it is possible to record video data to do off-line behavioural analysis. The web interface was created using PHP, JavaScript and CSS, which runs on an Apache web server on the Raspberry Pi and is based on Silvan Melchior’s implementation (https://github.com/silvanmelchior/RPi_Cam_Web_Interface). All the code we used and developed, is open-source and it can be downloaded from https://github.com/surjeets/Animal_Home_Cage and https://github.com/surjeets/Home-Cage-Python-and-MATLAB-Code.

The web interface has Home, Settings, Preview and GPIO Control tabs; by default, the opening page is the Home page (Fig. 1-E) where users can view the live camera feed, start/stop video recording, capture an image or stop the camera. Users can change the camera settings on the Settings page (e.g. video resolution, frame rate, brightness, contrast, sharpness, ISO, rotation, annotations, video splitting, etc.) (Fig. 2). Users can also preview/download/delete captured images and videos on the Preview page (Fig. 3). The storage space available on the SD card (or selected storage device) is indicated on this page. Further, users can also manually control external peripherals attached to the system (e.g. LEDs, buzzers, motors, etc.) via the GPIO Control interface (Fig. 4).

**Fig 2.**
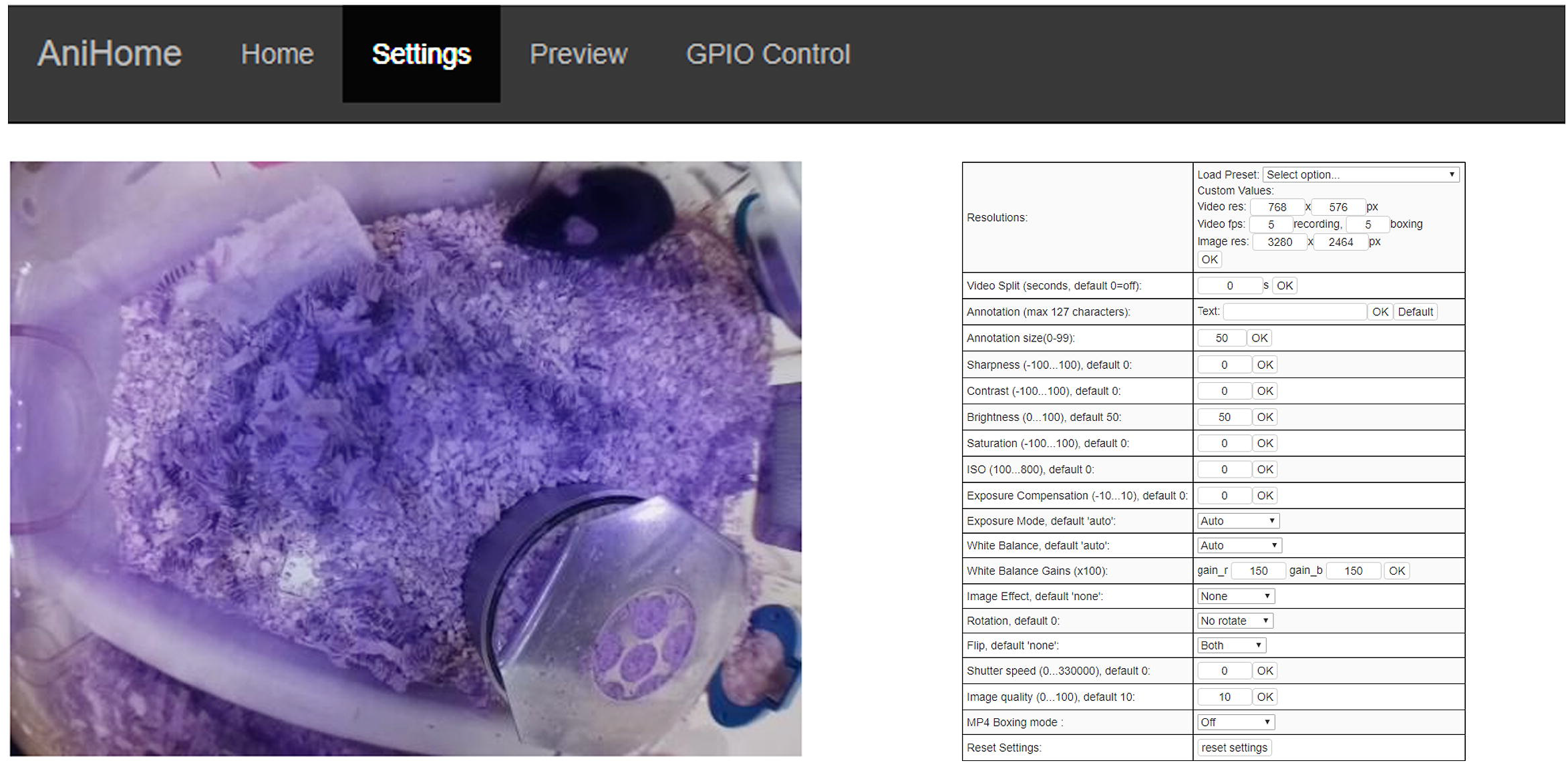
Web interface for camera settings. This interface allows a user to change camera settings such as: video resolution, frame rate, brightness, contrast, sharpness, ISO, rotation, annotations, video splitting etc.

**Fig 3.**
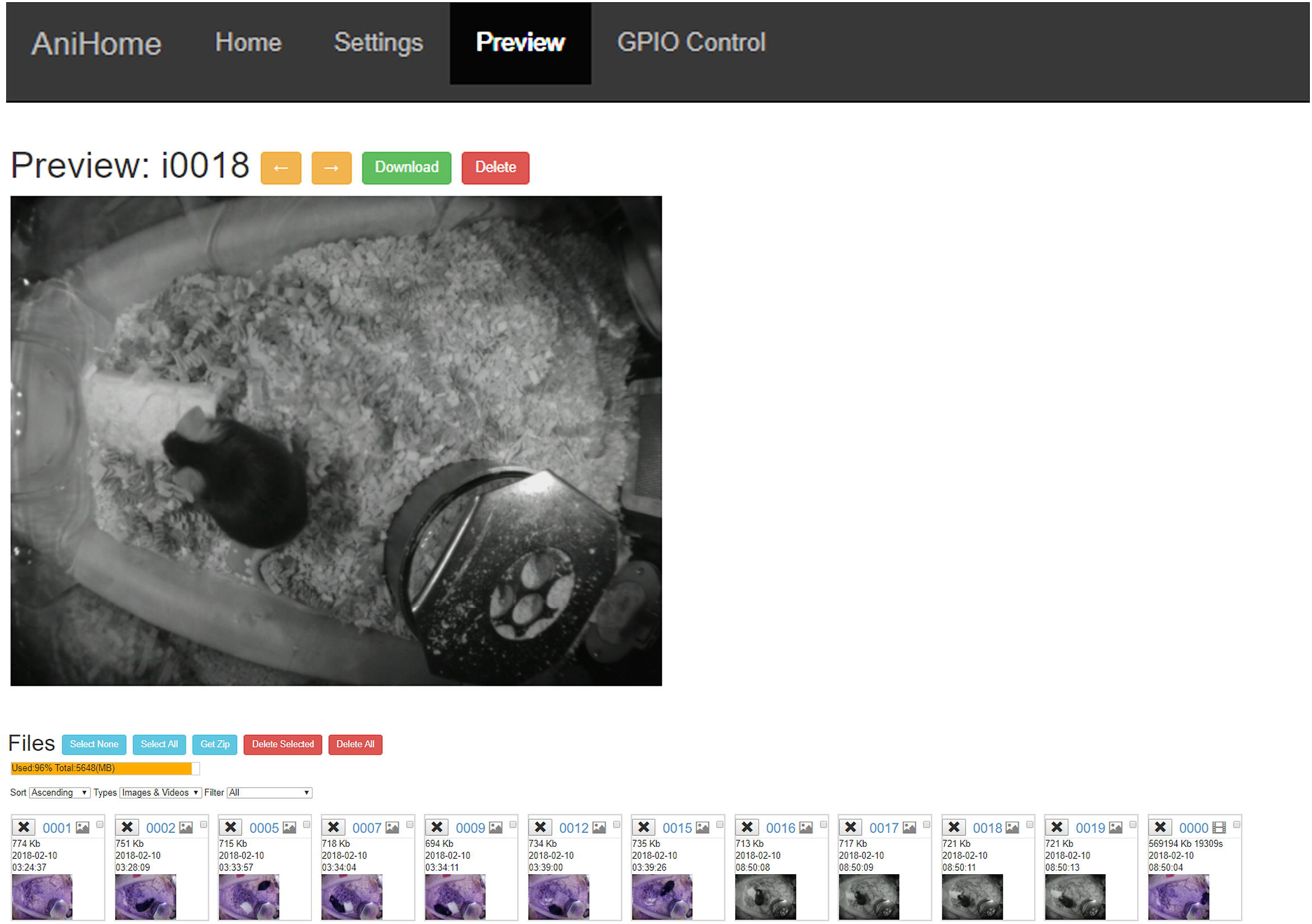
Preview of recorded videos/captured images. This interface allows the user to preview/download/delete captured images and videos. Further, storage space available on the SD card for recording more videos/capturing images is also indicated.

**Fig 4.**
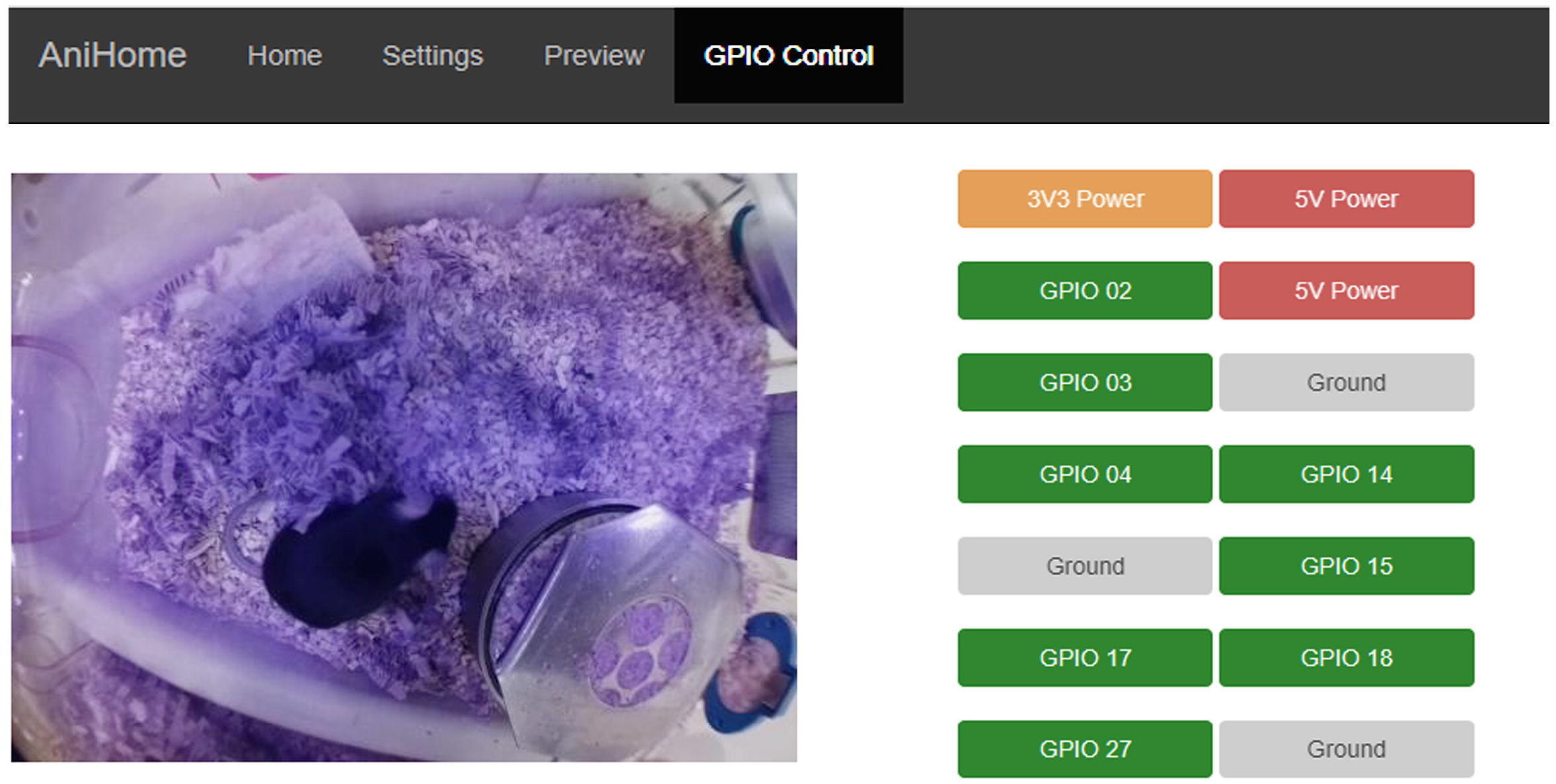
Remote control of external peripherals. This interface allows the user to manually control external peripherals attached to the system via the RPi GPIO pins (e.g. LEDs, Buzzers, motors, etc.).

The real-time video streaming is based on the program *RaspiMJPEG* which is an OpenMAX-application developed by Silvan Melchior and is based on the Multimedia Abstraction Layer (MMAL) library [33]. In the web interface of our system, the *RaspiMJPEG* makes a connection to the camera (MMAL) and generates a continuous stream of single frames which are saved in the /dev/shm/mjpeg directory of the RPi (e.g. cam.jpg). This folder is a virtual memory shared location that speeds up the execution by avoiding having to do many read/write to disk operations. These frames are accessed via URLs on the Apache Web server. The script cam_pic.php returns the latest one and cam_pic_new.php merges the frames into an mjpeg stream. The stream of the frames (preview images) is maintained throughout even during capturing/recording video/image. The captured video/image data is stored in the media folder (/media) of the RPi, the images are stored in JPEG format while the video is stored in h264 format but can be automatically packaged into an mp4 file when the recording ends. Alternatively, the video stream can also be assessed using the iSpy Connect software (www.ispyconnect.com) running on a windows machine by adding the following url: http://<your_pi_ip>:<port>/html/cam.jpg or http://<your_pi_ip>:<port>/html/cam_pic_new.php? in the JPEG URL or MJPEG URL tab within video source. This allows the user to simultaneously capture a video stream on a desktop computer and on the RPi. This implementation allows to schedule video recordings across multiple cages.

### Offline Behaviour Analysis

To process the recorded videos, we implemented two offline algorithms for behavioural data analysis using OpenCV and Python (https://github.com/surjeets/Home-Cage-Python-and-MATLAB-Code), one for motion detection and another for animal tracking in the home-cage. Using these simple algorithms, we were able to demonstrate our approach by performing sleep/wake analysis with accuracy comparable to EMG recordings based methods.

### Motion detection, tracking and sleep/wake scoring

We use a simple and well-known computer vision method to detect motion and to track animals in the home-cage. To estimate the amount of motion in a video we extract the background [34–37] and take the difference between consecutive frames (frame differencing) [38].

For motion detection we implemented the algorithm shown in Fig 5 (Algorithm 1). First, the background is estimated by calculating the weighted mean of previous frames along with the current frame. In this way our system can dynamically adjust to the background even with the small changes in the lighting conditions. After this, frame differencing is performed by subtracting the weighted mean from the current frame to get a delta frame:

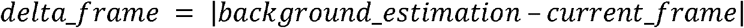

**Fig 5.**
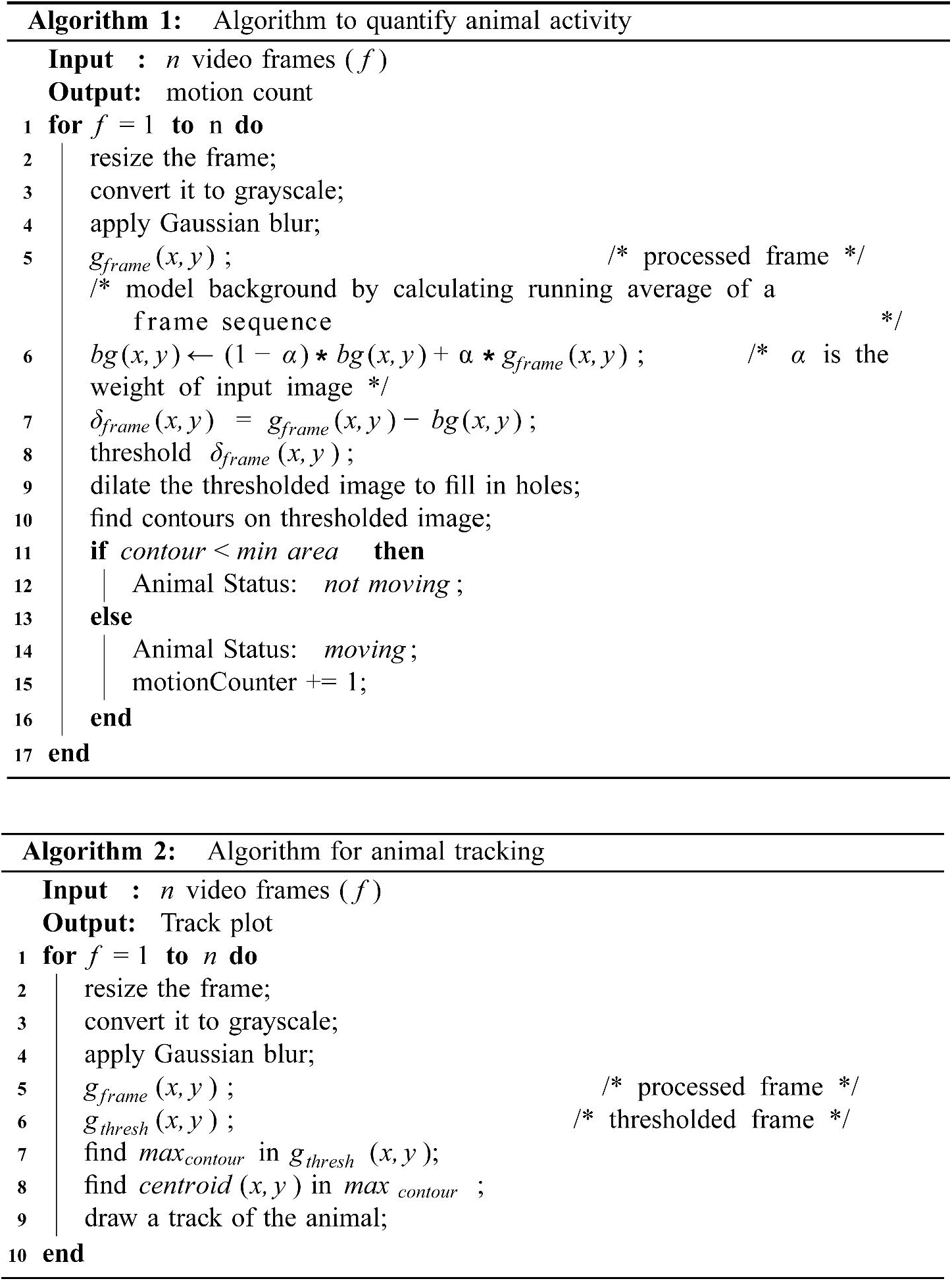
Off-line video analysis algorithms. Algorithm 1 (top) quantifies animal activity and Algorithm 2 (bottom) calculates and draws the trajectory of the animal detected. Both algorithms are presented in pseudo-code (in our repository, they are implemented in Python using openCV).

The delta frame is then thresholded (assigning a constant value to all the pixels above the threshold and zero to the rest) to find regions in the image that contain substantial difference from the background. These regions correspond to “motion regions” in each video frame.

Next, we calculated the area of each motion region in the thresholded delta frame. If these motion regions have an area larger than a threshold (the area that a mouse occupies in the visual field is assumed to be larger than a certain number of pixels), the motion is considered to be caused by the mouse (the motion region is shown with a rectangle around it Fig 6-A). This threshold was set to avoid other objects (e.g. cables or bedding) or pixel fluctuations due to breathing being incorrectly detected as mouse motion.

**Fig 6.**
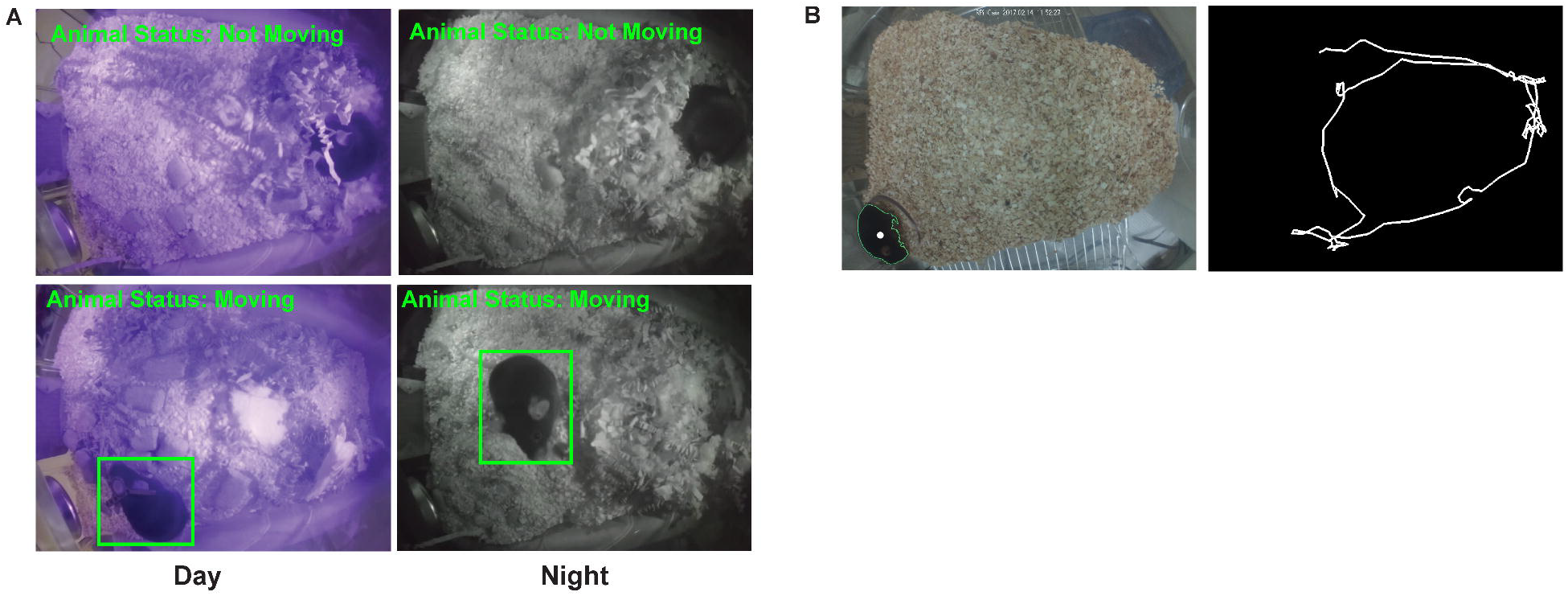
Animal motion detection and tracking. (A) Results of activity detection algorithm showing day (left column) and night (right column) time activity of the animal in the home-cage. (B) Results for animal tracking algorithm with the centroid marked with a white dot (left) and the tracking plot generated by connecting the detected centroids in each frame (right).

For animal tracking we implemented the algorithm shown in Fig 5 (Algorithm 2). This algorithm assumes that there is a high contrast between the mouse and the home-cage background. First, a Gaussian filter (size = 21, sigma = 3.5) is used to reduce noise in each video frame, then intensity-based thresholding is applied to segment the filtered image. Subsequently, to find the animal location, the contour with the largest area is detected, which, in the home-cage, is usually the animal. Finally, to calculate the trajectory of an animal, the centroids of the detected largest contour in each frame are connected (Fig 6-B). These trajectories can be used to calculate the distance travelled by the animal or to visualize its spatial occupancy.

We use the centroid of the motion-detected region to track the animal position in the home-cage (Fig 6B). These trajectories can be used to calculate the distance travelled by the animal or to visualize its spatial occupancy. Finally, we use our motion measure (motion signal) to predict when the animal is sleeping or awake. For sleep/wake scoring, we filtered the video-based motion signal below 0.1 Hz using a zero-lag Chebyshev filter and detrended using 10 sec windows with 5 sec overlap (using the locdetrend function from the Chronux Toolbox) to remove the variations in the baseline across light and dark cycle. The result was then rectified and integrated using 20-sec moving windows. Next, this signal was thresholded using its 50-60 percentile to classify waking and sleep periods. If the signal was below the threshold for at least 40 sec, the animal was assumed to be asleep, or awake otherwise. This threshold was chosen based on the optimal length based on the accuracy of sleep detection (Fig.9A). A similar criteria was reported in [27, 28, 39]. The EMG based sleep/wake scoring was performed by filtering the raw EMG activity between 90 Hz and 1000 Hz and then rectifying and integrating it using 20-sec moving windows. A state was scored as sleep when the EMG signal was smaller than its 50-60 percentile, otherwise it was classified as awake [27]. The accuracy of video-based classification was calculated by counting the amount of time the video-based scoring was the same as the EMG-based scoring divided by the total recording time:

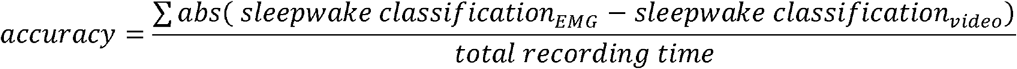

### Camera settings and temporal precision evaluation

In some behavioural tasks (e.g. reach and grasp, grooming behaviour) it is very important to acquire the data at high frame rates. With the Picamera camera board V2.1 (8MP Sony IMX219 sensor), it is possible to record video @ 30 fps in full resolution (no shutter lag and simultaneous recording), and also @120 fps if using the 2 × 2 analogue binning mode (which is a clocking scheme to collect the charge of a 2×2 CCD pixel area). We evaluated the temporal precision of the frame acquisition of the RPi Camera by comparing the average number of dropped frames and variation in interframe intervals (IFIV) in a video at different framerates (30, 60, 90 and 120 fps) and recording formats: h264, which is the compression standard MPEG-4 AVC and raw YUV, which is an uncompressed bitmap data format. In order to quantify the temporal precision of the frame acquisition of the Picamera, we recorded the time stamps of each frame acquisition into a csv file using the *picamera* python library [40].

To evaluate the time difference in simultaneous frame acquisition, multiple RPis were simultaneously triggered with an external square TTL pulse. On another RPi (besides the ones being triggered), we monitored the transition (high or low) of selected GPIO pins in real-time that corresponded to external trigger pulses that represent the start and stop times of video recording (Fig. 7) using *piscope* [41]. The timestamps of acquired frames on different RPis were later aligned offline to quantify any temporal differences in the acquisition. Table 1 shows that the jitter in the frame acquisition on different RPis is in the order of few microseconds.

**Table 1.** Comparison of frames per second (fps) and variance in Inter frame interval (IFIV) of the video data acquired over multiple trials: 15-minute video (320 X 240) was recorded in a raw (YUV) format and h264 encoding for 10 trials at a fixed frame rate (fps) on multiple RPis (n=5). ^*^results expressed as mean ± standard deviation.

| Expected frame rate | Frame rate achieved (raw YUV format)* | Variance in interframe interval (IFIV) for raw YUV format* | Frame rate achieved (h264 encoded)* | Variance in interframe interval (IFIV) for h264 encoded* |
| --- | --- | --- | --- | --- |
| 30 fps | $28.98 \pm 1.70$ fps | $39.70 \pm 66.83$ ms | $30.00 \pm 0.00$ fps | $2.00 \pm 0.18$ μs |
| 60 fps | $52.84 \pm 3.34$ fps | $81.34 \pm 27.66$ ms | $60.20 \pm 0.00$ fps | $0.63 \pm 0.31$ μs |
| 90 fps | $77.61 \pm 4.59$ fps | $54.45 \pm 31.78$ ms | $90.24 \pm 0.00$ fps | $2.23 \pm 0.35$ μs |
| 120 fps | N.A. | N.A. | $120.54 \pm 0.00$ fps | $2.44 \pm 0.46$ μs |

**Fig 7.**
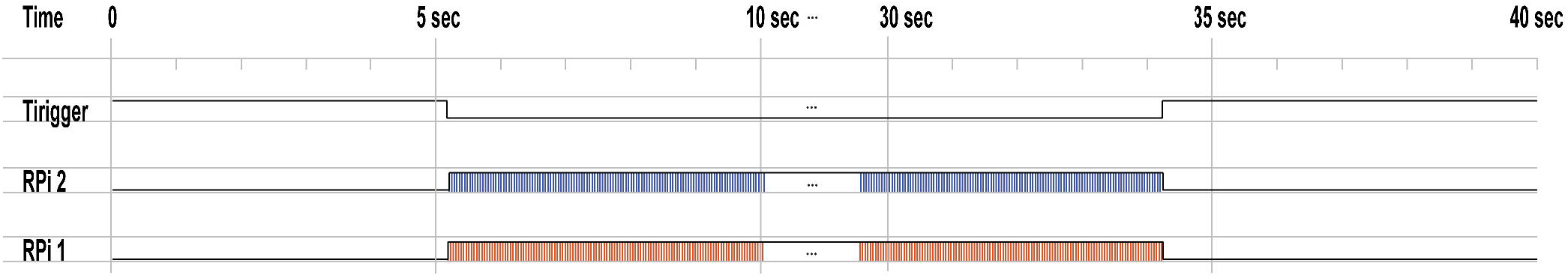
Simultaneous video acquisition on multiple Raspberry Pis. Example of two Raspberry Pis (RPi) triggered by an external TTL pulse (Trigger) to simultaneously start/stop video recording RPis (RPi 1 and RPi2).

### Off-line alignment of the video with additional signals

To align the video recordings to LFP or EMG signals, we keep track of the times when the video acquisition begins and ends in the same acquisition system that records the electrophysiological signals. We do this by setting a TTL signal (Transistor-Transistor Logic 5V) to HIGH in one of the GPIO pins of the RPi when the video acquisition starts and set to LOW when the acquisition terminates. Additionally, we also record the times when each video frame is acquired (see https://picamera.readthedocs.io/en/release-1.13/api_camera.html).

## Results

### High Frame Rate Acquisition

The temporal precision and resolution required for the video recordings depends largely on the behaviour we are interested in. For example, if one is interested in skilled reaching or play behaviour this might require to record video at higher resolution and frame rates than if we are interested in monitoring the general activity in the home-cage. Since the temporal precision of the video varies depending on the settings we used, we compared some of the file formats and recording settings in this system. We observed that there are dropped frames when we save the video in raw YUV format above 30 fps at full resolution. This situation gets worse when the frame rate augments. For example, at 90 fps we can only achieve a frame rate of 77.61 ± 4.59 fps with a lag of 54.45 ± 31.78 ms between consecutive frames (Table 1). In contrast, using the H264 encoding we were able to record at 120 fps with no missed frames (120.54 ± 0.00 fps) and almost no lag between frames (2.44 ± 0.46 µs).

### Multiple camera synchronization

The system presented here assumes that a single camera is placed on the home-cage providing a top-down view. However, it is possible to use more than one camera if they can be synchronized [42]. To test this, we evaluated the temporal alignment of two cameras recording video synchronously (see Methods). We found that the temporal difference between the two cameras is in the order of less than ten microseconds when using H264 encoding at 320 x 240 pixel resolution and frame rates from 30-120 (last column of Table 1 and Fig. 7).

### Sleep-wake state classification

To classify the behaviour of the mouse into sleep and wake states, we applied the following criteria: if the mouse was not moving (according to our motion detection algorithm) for more than 40 sec, we assumed that it was asleep, otherwise we assumed it was awake (see Methods and Fig 5). To validate this classification, we simultaneously recorded hippocampal local field potentials (LFP) and EMG signals and compared them to our video-only based classification (Fig. 8A-B). The high-throughput methodology for sleep-wake monitoring presented here can be easily incorporated into any existing phenotyping test battery, e.g. circadian studies (Fig. 8B). For example, our 24 hour sleep/wake classification algorithm shows that animals were more active during the dark cycle than during the light cycle (Fig.8C-D, n=5, one-way anova, Bonferroni correction, F_(1,9)_ = 12.03, p = 0.0085).

**Fig 8.**
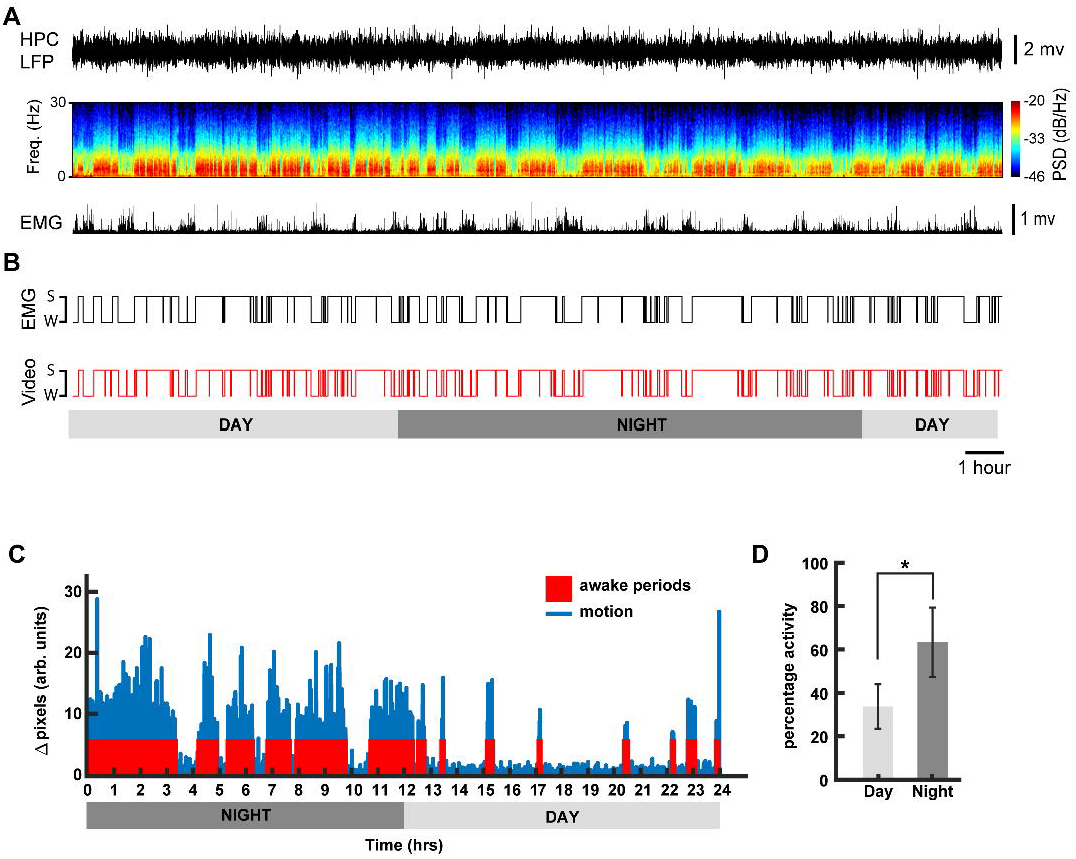
Sleep-wake state classification. (A) Example of hippocampal (HPC) LFP and its spectrogram and rectified EMG activity recorded from a naturally sleeping animal during 24h (B) Comparison of hypnogram scoring the sleep and waking states calculated based on the EMG and video monitoring. (C) Example actogram generated using Algorithm 1, for a day-night sleep-wake state classification for one mouse. (D) Mean estimated awake duration expressed in terms of percentage activity for day and night cycles (12 hr each) for n = 5 animals (mean ± standard deviation, ^*^ denotes p < 0.01; one-way ANOVA using Bonferroni post hoc).

Our sleep-wake classification was highly correlated to the 24 hr LFP/EMG recordings (90.52% ± 0.4 accuracy. Fig. 9C). Similar results were obtained for shorter recordings (data not shown).

**Fig. 9.**
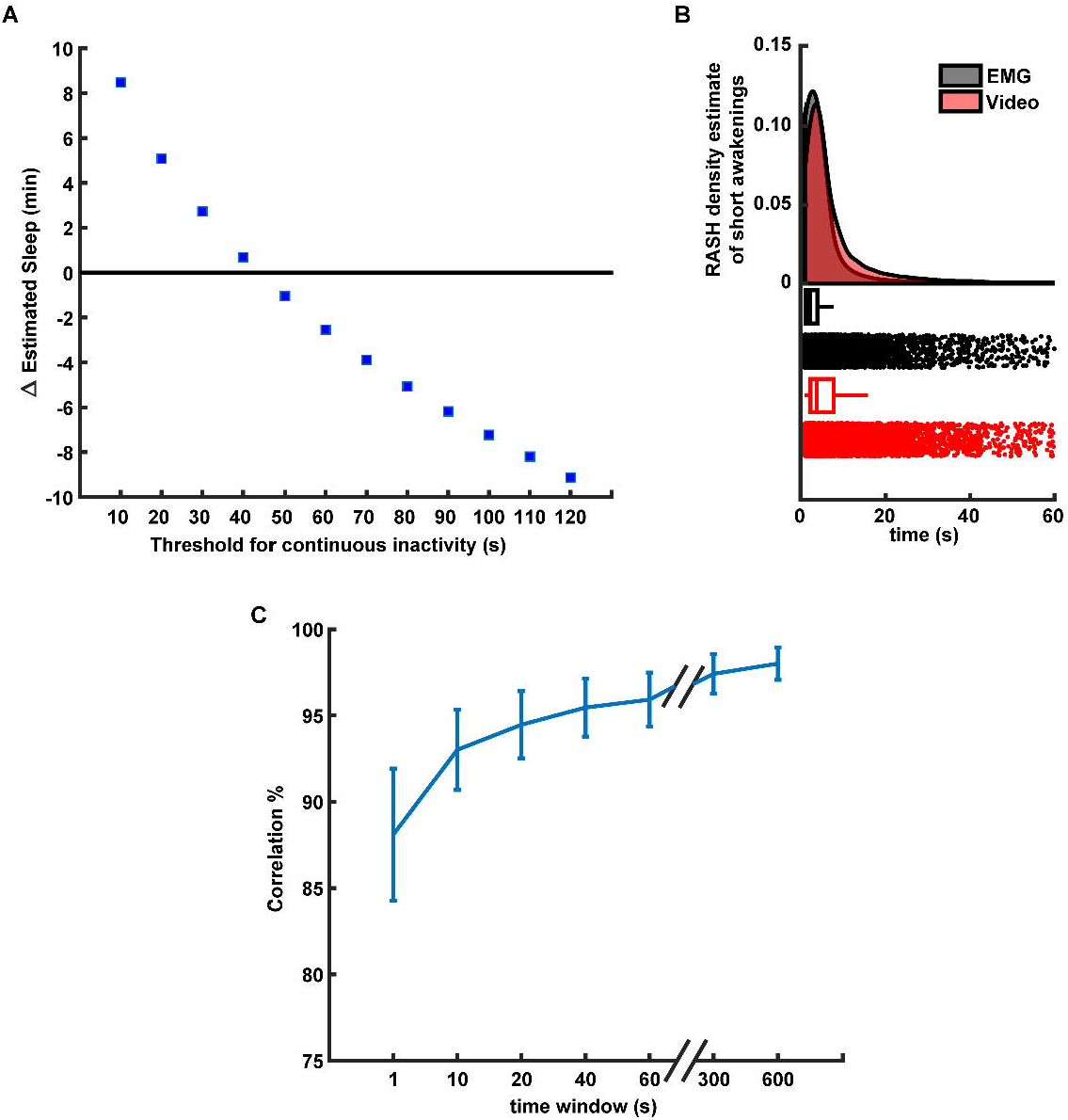
Performance of video-based sleep/wake classification. (A) Relationship between the time length of continuous inactivity threshold and accuracy of sleep/wake video-based classification. Y-axis shows the difference in minutes between the average amounts of sleep detected using the video-based and EMG classification in 2 hr interval (n=5 animals). (B) Density estimations of sleep events durations for EMG-based (gray) and video-based (red) classification (top). The density was estimated using the RASH method described in [43]. Box and raster plots (bottom) of the sleep detected events of the different durations in seconds (X-axis). (C) Correlation between the video-based and EMG-based sleep/wake classifications for different smoothing windows durations. Error bars represent the S.E.M for n=5 animals.

We chose a 40 sec threshold to determine the behavioural state of the animal based on the optimal accuracy of our method compared to the EMG based classification (Fig. 9A). A very similar finding is reported in [28]. The performance of our method decreased when using shorter windows thresholds because the mouse could be immobile for very short periods of time but not asleep. Analogously, when using long windows, the accuracy drops because mice can sleep during short periods (see Discussion). We found that the estimated distributions of the duration of the detected sleep events using the video-based and EMG based methods is generally similar. However, they differ when the durations of the events are small, close to 1-2 sec or between 10-30 seconds (Fig. 9B). In addition, the accuracy of the video-based method changes with respect to the noise in the motion detection signal, which in turn is modulated (in part) by the smoothening of this signal (see Methods). When the video-based motion detection signal was smoothened using very short windows, small motion fluctuations could be incorrectly detected as wakening. The smoothening over longer windows improves the accuracy of the method up until an asymptotic value (Fig. 9C).

## Discussion

When studying complex behaviour, experimental designs that require the experimenter to move the animal from its home-cage environment to an unfamiliar apparatus (novel environment) are relatively disruptive, time- and labour-consuming and require additional laboratory space. This disruption may influence behaviour, general well-being, and metabolism of the animal, affecting the phenotypic outcome, even if the data collection method is automated, therein possibly creating spurious findings. Therefore, monitoring the animals in their home-cage has several advantages. Though there are already open-source and commercial solutions available to monitor animal behaviour [44–51], they are usually difficult to construct, require a new infrastructure/space, and are expensive. Conversely, we propose a video-based system that is composed of off-the-shelf components and designed to be simple and easily integrated in existing home-cage setups with minimal modifications for remote behaviour monitoring. Such a system can be used for short-term welfare assessment (e.g. post-surgery monitoring) by enabling 24 hr monitoring, even in the dark phase where welfare assessment without disturbance to the cage is difficult and subjective. Further, video data collected with this system can be used to classify sleep-wake states of an animal in the home-cage using video-based tracking. The simple video-based tracking algorithm that we use has an accuracy above 90% when compared with tethered LFP/EMG recordings for sleep-wake state classification. This algorithm is based on the quantification of motion during a period to determine if the animal is awake or asleep. We found that the highest accuracy of the sleep/wake classification is reached when the time window threshold for no motion period to be considered as sleep is close to 40 secs compared to the gold standard EEG/EMG based classification. The performance of our classification method mainly depends on two factors: the amount of time we use as a threshold to assume the animal is asleep and the amount of noise (or smoothness) in the motion detection signal. When considering shorter time windows as a threshold, this method incorrectly classifies periods of quiet wakefulness as sleep. Analogously, when the window is too large, this method incorrectly classifies long periods of sleep that contain small movement as wake periods (e.g. twitches) (Fig. 9A). The smoothness of the motion detection signal is determined (in part) by the length of the smoothening window. If the window is small, the motion detection signal contains small changes in the home-cage (illumination, cables, bedding, etc), that can cause the algorithm to incorrectly detect these changes as the mouse being awake. We can alleviate this problem by extending the smoothening signal to reach an asymptotic performance compared to the EMG-based classification (Fig. 9C). This has been previously reported in similar video-based sleep/wake classification approaches [27, 28]. Although these results show that our video-based algorithm reliably determines whether the animal is awake or asleep (compared to an LFP-EMG based algorithm), for a more in depth classification of sleep into Rapid-Eye Movement (REM) and non-Rapid Eye Movement (NREM) sleep states, our video analysis should be complemented with more advance computer vision techniques [52] or additional brain signals.

Our system has some advantages and limitations due to the technology we use. For example, an advantage of the video based tracking algorithm presented here it is a higher temporal resolution (in the order of milliseconds) and spatial resolution (in the order of millimeters), both determined by the video camera, as compared to standard tracking approaches such as RFID systems. In contrast, a home-cage RFID system with an array of 18 RFID sensors (ID20LA) has a spatial resolution of 5 cm and a temporal resolution of 1Hz [24]. One disadvantage of our system is that it can only track one animal at a time due to the potential high similarity between multiple mice (e.g. same fur colour and shape) [47], compared to the multiple animal tracking capability of an RFID based system. However, RFID (and video-hybrid) systems are highly specialized, difficult to setup and expensive [46, 51, 53, 54]. A potential approach to improve video-based tracking methods is to extract high level features from each animal to identify every individual using machine learning [55, 56]. However, such algorithms need to be highly calibrated and tested to provide reliable results. Another possible limitation of our system is the processing power of the RPi if extensive online video processing is required. For example, streaming live video over a network and simultaneously performing object recognition on the RPi in real time might result in eventual frame dropping. Despite these limitations, the system has several advantages for large scale monitoring of rodents: it can be easily integrated with any available rodent cage-rack system. Further, a user can remotely monitor day/night activity of animals in multiple cages, from virtually anywhere via the web interface which can be applied not only for research but animal care and husbandry.

Finally, in addition to the sleep/wake classification application presented in this work, our system could also be applied to study different aspects of animal behaviour. For example, our system can produce actograms and determine the proportion of sleep and wake periods during the day (Fig. 8C-D). A similar trend is observed in systems using running wheels for phenotyping circadian rhythms [39]. Even though running wheels are widely used to study circadian rhythm in rodents, it has been shown that long-term access to a running wheel affect animal behaviour and disease pathology [26]. The home-cage video monitoring system does not introduce such confounding factors, rather it allows researchers to evaluate additional behavioural parameters such as the distance travelled, or the time spent in certain areas of the cage, which can provide additional data on anxiety or behavioural inhibition.

## Conclusions

In this paper, we present a low-cost home-cage reliable behavioural monitoring system that can be easily adapted and extended to a large variety of conditions and pre-existing experimental setups. We demonstrated that our system can be used to study natural sleep patterns of mice in their home-cage using only video. Due to its suitability for remote behaviour monitoring and minimization of experimenter bias at low-cost, systems like the one presented here, can become a useful tool for high-throughput data collection and behavioural analysis of rodents in their home-cage.

## Acknowledgements

This work was supported by Natural Sciences and Engineering Research Council of Canada (NSERC) Discovery Grant #40352 and #RGPIN-2017-03857 (MHM), Campus Alberta for Innovation Program Chair, Alberta Alzheimer Research Program, Alzheimer Society of Canada (MHM), NSERC Discovery Grant and Alberta Neuroscience Program Grant (RJS).

## Author contributions

S.S., E.B.C., and M.H.M. conceived and designed the method, S.S., E.B.C. and M.N. performed experimental work, S.S. and M.N. performed data analysis, S.S. and E.B.C wrote the manuscript, which all authors commented on and edited. M.H.M. and R.J.S. provided project leadership.

## References

1. Dombeck DA, Khabbaz AN, Collman F, Adelman TL, Tank DW. Imaging large-scale neural activity with cellular resolution in awake, mobile mice. Neuron. 2007;56(1):43–57. Epub 2007/10/09. doi: 10.1016/j.neuron.2007.08.003. PubMed PMID: 17920014; PubMed Central PMCID: PMCPMC2268027.

2. Ghosh KK, Burns LD, Cocker ED, Nimmerjahn A, Ziv Y, Gamal AE, et al. Miniaturized integration of a fluorescence microscope. Nat Methods. 2011;8(10):871–8. Epub 2011/09/13. doi: 10.1038/nmeth.1694. PubMed PMID: 21909102; PubMed Central PMCID: PMCPMC3810311.

3. Grinvald A, Hildesheim R. VSDI: a new era in functional imaging of cortical dynamics. Nat Rev Neurosci. 2004;5(11):874–85. Epub 2004/10/22. doi: 10.1038/nrn1536. PubMed PMID: 15496865.

4. Luo L, Callaway EM, Svoboda K. Genetic dissection of neural circuits. Neuron. 2008;57(5):634–60. Epub 2008/03/18. doi: 10.1016/j.neuron.2008.01.002. PubMed PMID: 18341986; PubMed Central PMCID: PMCPMC2628815.

5. Mohajerani MH, Chan AW, Mohsenvand M, LeDue J, Liu R, McVea DA, et al. Spontaneous cortical activity alternates between motifs defined by regional axonal projections. Nat Neurosci. 2013;16(10):1426–35. Epub 2013/08/27. doi: 10.1038/nn.3499. PubMed PMID: 23974708; PubMed Central PMCID: PMCPMC3928052.

6. Zhang F, Aravanis AM, Adamantidis A, de Lecea L, Deisseroth K. Circuit-breakers: optical technologies for probing neural signals and systems. Nat Rev Neurosci. 2007;8(8):577–81. Epub 2007/07/24. doi: 10.1038/nrn2192. PubMed PMID: 17643087.

7. Bermudez-Contreras E, Chekhov S, Sun J, Tarnowsky J, McNaughton BL, Mohajerani MH. High-performance, inexpensive setup for simultaneous multisite recording of electrophysiological signals and mesoscale voltage imaging in the mouse cortex. Neurophotonics. 2018;5(2):025005. Epub 2018/04/14. doi: 10.1117/1.NPh.5.2.025005. PubMed PMID: 29651448; PubMed Central PMCID: PMCPMC5874445.

8. Saito T, Matsuba Y, Mihira N, Takano J, Nilsson P, Itohara S, et al. Single App knock-in mouse models of Alzheimer’s disease. Nat Neurosci. 2014;17(5):661–3. Epub 2014/04/15. doi: 10.1038/nn.3697. PubMed PMID: 24728269.

9. Janus C, Westaway D. Transgenic mouse models of Alzheimer’s disease. Physiol Behav. 2001;73(5):873–86. Epub 2001/09/22. PubMed PMID: 11566220.

10. Elder GA, Gama Sosa MA, De Gasperi R. Transgenic mouse models of Alzheimer’s disease. Mt Sinai J Med. 2010;77(1):69–81. Epub 2010/01/27. doi: 10.1002/msj.20159. PubMed PMID: 20101721; PubMed Central PMCID: PMCPMC2925685.

11. Singh S, Kaur H, Sandhir R. Fractal dimensions: A new paradigm to assess spatial memory and learning using Morris water maze. Behav Brain Res. 2016;299:141–6. Epub 2015/11/26. doi: 10.1016/j.bbr.2015.11.023. PubMed PMID: 26592165.

12. Goodman S, Check E. The great primate debate. Nature. 2002;417(6890):684–7. Epub 2002/06/18. doi: 10.1038/417684a. PubMed PMID: 12066153.

13. Spruijt BM, DeVisser L. Advanced behavioural screening: automated home cage ethology. Drug Discov Today Technol. 2006;3(2):231–7. Epub 2006/07/01. doi: 10.1016/j.ddtec.2006.06.010. PubMed PMID: 24980412.

14. Howerton CL, Garner JP, Mench JA. A system utilizing radio frequency identification (RFID) technology to monitor individual rodent behavior in complex social settings. J Neurosci Methods. 2012;209(1):74–8. Epub 2012/06/16. doi: 10.1016/j.jneumeth.2012.06.001. PubMed PMID: 22698663.

15. Steele AD, Jackson WS, King OD, Lindquist S. The power of automated high-resolution behavior analysis revealed by its application to mouse models of Huntington’s and prion diseases. Proc Natl A cad Sci U S A. 2007;104(6):1983–8. Epub 2007/01/31. doi: 10.1073/pnas.0610779104. PubMed PMID: 17261803; PubMed Central PMCID: PMCPMC1794260.

16. de Visser L, van den Bos R, Kuurman WW, Kas MJ, Spruijt BM. Novel approach to the behavioural characterization of inbred mice: automated home cage observations. Genes Brain Behav. 2006;5(6):458–66. Epub 2006/08/23. doi: 10.1111/j.1601-183X.2005.00181.x. PubMed PMID: 16923150.

17. Tecott LH, Nestler EJ. Neurobehavioral assessment in the information age. Nat Neurosci. 2004;7(5):462–6. Epub 2004/04/29. doi: 10.1038/nn1225. PubMed PMID: 15114359.

18. Winter Y, Schaefers AT. A sorting system with automated gates permits individual operant experiments with mice from a social home cage. J Neurosci Methods. 2011;196(2):276–80. Epub 2011/01/25. doi: 10.1016/j.jneumeth.2011.01.017. PubMed PMID: 21256865.

19. Hawkins P. Recognizing and assessing pain, suffering and distress in laboratory animals: a survey of current practice in the UK with recommendations. Lab Anim. 2002;36(4):378–95. Epub 2002/10/25. doi: 10.1258/002367702320389044. PubMed PMID: 12396281.

20. Littin K, Acevedo A, Browne W, Edgar J, Mendl M, Owen D, et al. Towards humane end points: behavioural changes precede clinical signs of disease in a Huntington’s disease model. Proc Biol Sci. 2008;275(1645):1865–74. Epub 2008/05/08. doi: 10.1098/rspb.2008.0388. PubMed PMID: 18460434; PubMed Central PMCID: PMCPMC2593928.

21. Storch C, Hohne A, Holsboer F, Ohl F. Activity patterns as a correlate for sleep-wake behaviour in mice. Journal of Neuroscience Methods. 2004;133(1-2):173–9. doi: 10.1016/j.jneumeth.2003.10.008. PubMed PMID: WOS:000188887700020.

22. Flores AE, Flores JE, Deshpande H, Picazo JA, Xie XMS, Franken P, et al. Pattern recognition of sleep in rodents using piezoelectric signals generated by gross body movements. Ieee T Bio-Med Eng. 2007;54(2):225–33. doi: 10.1109/Tbme.2006.886938. PubMed PMID: WOS:000243952100004.

23. Brown L, Hasan S, Foster R, Peirson S. COMPASS: Continuous Open Mouse Phenotyping of Activity and Sleep Status [version 1; referees: 3 approved, 1 approved with reservations] 2016.

24. Noorshams O. Automated systems to assess weights and activity in group housed mice. Vancouver, BC: University of British Columbia; 2017.

25. Chemelli RM, Willie JT, Sinton CM, Elmquist JK, Scammell T, Lee C, et al. Narcolepsy in orexin knockout mice: Molecular genetics of sleep regulation. Cell. 1999;98(4):437–51. doi: Doi 10.1016/S0092-8674(00)81973-X. PubMed PMID: WOS:000082174900005.

26. Richter H, Ambree O, Lewejohann L, Herring A, Keyvani K, Paulus W, et al. Wheel-running in a transgenic mouse model of Alzheimer’s disease: protection or symptom? Behav Brain Res. 2008;190(1):74–84. Epub 2008/03/18. doi: 10.1016/j.bbr.2008.02.005. PubMed PMID: 18342378.

27. Fisher SP, Godinho SIH, Pothecary CA, Hankins MW, Foster RG, Peirson SN. Rapid Assessment of Sleep-Wake Behavior in Mice. J Biol Rhythm. 2012;27(1):48–58. doi: 10.1177/0748730411431550. PubMed PMID: WOS:000299848100005.

28. Pack AI, Galante RJ, Maislin G, Cater J, Metaxas D, Lu S, et al. Novel method for high-throughput phenotyping of sleep in mice. Physiol Genomics. 2007;28(2):232–8. doi: 10.1152/physiogenomics.00139.2006. PubMed PMID: WOS:000244171700010.

29. Jafari Z, Mehla J, Kolb BE, Mohajerani MH. Prenatal noise stress impairs HPA axis and cognitive performance in mice. Sci Rep. 2017;7(1):10560. Epub 2017/09/07. doi: 10.1038/s41598-017-09799-6. PubMed PMID: 28874680; PubMed Central PMCID: PMCPMC5585382.

30. Mehla J, Lacoursiere S, Stuart E, McDonald RJ, Mohajerani MH. Gradual Cerebral Hypoperfusion Impairs Fear Conditioning and Object Recognition Learning and Memory in Mice: Potential Roles of Neurodegeneration and Cholinergic Dysfunction. J Alzheimers Dis. 2018;61(1):283–93. Epub 2017/11/21. doi: 10.3233/JAD-170635. PubMed PMID: 29154281.

31. Lee JQ, Sutherland RJ, McDonald RJ. Hippocampal damage causes retrograde but not anterograde memory loss for context fear discrimination in rats. Hippocampus. 2017;27(9):951–8. Epub 2017/07/08. doi: 10.1002/hipo.22759. PubMed PMID: 28686806.

32. Whishaw IQ, Faraji J, Kuntz J, Mirza Agha B, Patel M, Metz GAS, et al. Organization of the reach and grasp in head-fixed vs freely-moving mice provides support for multiple motor channel theory of neocortical organization. Exp Brain Res. 2017;235(6):1919–32. Epub 2017/03/21. doi: 10.1007/s00221-017-4925-4. PubMed PMID: 28315945.

33. Melchior S. RaspiCam Documentation. 2015.

34. KaewTraKulPong P, Bowden R. An Improved Adaptive Background Mixture Model for Real-time Tracking with Shadow Detection. In: Remagnino P, Jones GA, Paragios N, Regazzoni CS, editors. Video-Based Surveillance Systems: Computer Vision and Distributed Processing. Boston, MA: Springer US; 2002. p. 135–44.

35. Zivkovic Z, editor Improved adaptive Gaussian mixture model for background subtraction. Proceedings of the 17th International Conference on Pattern Recognition, 2004 ICPR 2004; 2004 23–26 Aug. 2004.

36. Zivkovic Z, Heijden Fvd. Efficient adaptive density estimation per image pixel for the task of background subtraction. Pattern Recogn Lett. 2006;27(7):773–80. doi: 10.1016/j.patrec.2005.11.005.

37. Godbehere AB, Matsukawa A, Goldberg K, editors. Visual tracking of human visitors under variable-lighting conditions for a responsive audio art installation. 2012 American Control Conference (ACC); 2012 27–29 June 2012.

38. Himani S. Parekh DGT, Udesang K. Jaliy. A Survey on Object Detection and Tracking Methods. International Journal of Innovative Research in Computer and Communication Engineering. 2014;2(2).

39. Eckel-Mahan K, Sassone-Corsi P. Phenotyping Circadian Rhythms in Mice. Curr Protoc Mouse Biol. 2015;5(3):271–81. Epub 2015/09/04. doi: 10.1002/9780470942390.mo140229. PubMed PMID: 26331760; PubMed Central PMCID: PMCPMC4732881.

40. Jones D. picamera a pure Python interface to the Raspberry Pi camera module. 2017.

41. Joan. Piscope is a digital waveform viewer for the Raspberry. 2017.

42. Salem GH, Dennis JU, Krynitsky J, Garmendia-Cedillos M, Swaroop K, Malley JD, et al. SCORHE: a novel and practical approach to video monitoring of laboratory mice housed in vivarium cage racks. Behav Res Methods. 2015;47(1):235–50. Epub 2014/04/08. doi: 10.3758/s13428-014-0451-5. PubMed PMID: 24706080; PubMed Central PMCID: PMCPMC4570574.

43. Bourel M, Fraiman R, Ghattas B. Random average shifted histograms. Comput Stat Data An. 2014;79:149–64. doi: 10.1016/j.csda.2014.05.004. PubMed PMID: WOS:000340139900011.

44. Bains RS, Wells S, Sillito RR, Armstrong JD, Cater HL, Banks G, et al. Assessing mouse behaviour throughout the light/dark cycle using automated in-cage analysis tools. Journal of Neuroscience Methods. 2017. doi: https://doi.org/10.1016/j.jneumeth.2017.04.014.

45. Jhuang H, Garrote E, Mutch J, Yu X, Khilnani V, Poggio T, et al. Automated home-cage behavioural phenotyping of mice. Nat Commun. 2010;1:68. Epub 2010/09/16. doi: 10.1038/ncomms1064. PubMed PMID: 20842193.

46. Weissbrod A, Shapiro A, Vasserman G, Edry L, Dayan M, Yitzhaky A, et al. Automated long-term tracking and social behavioural phenotyping of animal colonies within a semi-natural environment. Nat Commun. 2013;4:2018. Epub 2013/06/19. doi: 10.1038/ncomms3018. PubMed PMID: 23771126.

47. Hong W, Kennedy A, Burgos-Artizzu XP, Zelikowsky M, Navonne SG, Perona P, et al. Automated measurement of mouse social behaviors using depth sensing, video tracking, and machine learning. Proc Natl Acad Sci U S A. 2015;112(38):E5351–60. Epub 2015/09/12. doi: 10.1073/pnas.1515982112. PubMed PMID: 26354123; PubMed Central PMCID: PMCPMC4586844.

48. Aoki R, Tsubota T, Goya Y, Benucci A. An automated platform for high-throughput mouse behavior and physiology with voluntary head-fixation. Nat Commun. 2017;8(1):1196. Epub 2017/11/01. doi: 10.1038/s41467-017-01371-0. PubMed PMID: 29084948; PubMed Central PMCID: PMCPMC5662625.

49. Poddar R, Kawai R, Olveczky BP. A fully automated high-throughput training system for rodents. PLoS One. 2013;8(12):e83171. Epub 2013/12/19. doi: 10.1371/journal.pone.0083171. PubMed PMID: 24349451; PubMed Central PMCID: PMCPMC3857823.

50. Murphy TH, Boyd JD, Bolanos F, Vanni MP, Silasi G, Haupt D, et al. High-throughput automated home-cage mesoscopic functional imaging of mouse cortex. Nat Commun. 2016;7:11611. Epub 2016/06/14. doi: 10.1038/ncomms11611. PubMed PMID: 27291514; PubMed Central PMCID: PMCPMC4909937.

51. Bains RS, Cater HL, Sillito RR, Chartsias A, Sneddon D, Concas D, et al. Analysis of Individual Mouse Activity in Group Housed Animals of Different Inbred Strains using a Novel Automated Home Cage Analysis System. Frontiers in Behavioral Neuroscience. 2016;10(106). doi: 10.3389/fnbeh.2016.00106.

52. McShane BB, Galante RJ, Biber M, Jensen ST, Wyner AJ, Pack AI. Assessing REM sleep in mice using video data. Sleep. 2012;35(3):433–42. Epub 2012/03/02. doi: 10.5665/sleep.1712. PubMed PMID: 22379250; PubMed Central PMCID: PMCPMC3274345.

53. Redfern WS, Tse K, Grant C, Keerie A, Simpson DJ, Pedersen JC, et al. Automated recording of home cage activity and temperature of individual rats housed in social groups: The Rodent Big Brother project. PLoS One. 2017;12(9):e0181068. Epub 2017/09/07. doi: 10.1371/journal.pone.0181068. PubMed PMID: 28877172; PubMed Central PMCID: PMCPMC5587114.

54. Dandan S, Lin X, editors. A Hybrid Video and RFID Tracking System for Multiple Mice in Lab Environment. 2016 3rd International Conference on Information Science and Control Engineering (ICISCE); 2016 8–10 July 2016.

55. Perez-Escudero A, Vicente-Page J, Hinz RC, Arganda S, de Polavieja GG. idTracker: tracking individuals in a group by automatic identification of unmarked animals. Nat Methods. 2014;11(7):743–8. Epub 2014/06/02. doi: 10.1038/nmeth.2994. PubMed PMID: 24880877.

56. Mathis A, Mamidanna P, Cury KM, Abe T, Murthy VN, Mathis MW, et al. DeepLabCut: markerless pose estimation of user-defined body parts with deep learning. Nat Neurosci. 2018;21(9):1281–9. Epub 2018/08/22. doi: 10.1038/s41593-018-0209-y. PubMed PMID: 30127430.

